# Mediator-29 Limits *Caenorhabditis elegans* Fecundity

**DOI:** 10.1101/2025.02.14.638312

**Authors:** Qi Fan, Christopher Tran, Wei Cao, Roger Pocock

## Abstract

Mediator is an evolutionarily conserved multiprotein complex that acts as a critical coregulator of RNA polymerase II-mediated transcription. While core Mediator components are broadly required for transcription, others govern specific regulatory modules and signalling pathways. Here, we investigated the function of MDT-29/MED29 in the *Caenorhabditis elegans* germ line. We found that endogenously-tagged MDT-29 is ubiquitously expressed and concentrated in discrete foci within germ cell nuclei. Functionally, depleting MDT-29 in the germ line during larval development boosted fecundity. We determined that the increase in progeny production was likely caused by a combination of an expanded germline stem cell pool and decreased germ cell apoptosis. Thus, MDT-29 may act to optimize specific gene expression programs to control distinct germ cell behaviors, providing flexibility to progeny production in certain environments.

## INTRODUCTION

Faithful gamete production requires coordinated regulation of germ cell development and behavior. To this end, specific gene regulatory networks control germ cell proliferation (self-renewal), differentiation, survival, and sex determination. These regulatory mechanisms provide robustness to germ cell trajectories but also flexibility under certain environmental and physiological states. Thus, manipulation of specific gene regulatory networks and developmental decisions can affect fecundity.

Mediator is a multi-subunit complex with numerous roles in controlling transcription, including bridging interactions between RNA polymerase II and transcription factors (TFs) (Allen and Taatjes, 2015). Mediator contains ∼30 protein subunits that are grouped into discrete segments: head, middle, tail, and kinase modules (Figure 1A). In general, the head and middle modules coordinate assembly of the transcriptional pre-initiation complex and general recruitment to promoters; whereas the tail and kinase modules facilitate interactions with specific TFs and enhancer elements to regulate specialized developmental programs (Allen and Taatjes, 2015). As a result, selectively removing or including tail and kinase module components may give gene regulatory networks flexibility in controlling certain cellular events. Such functional specificity is exemplified by *Caenorhabditis elegans* tail module subunits MDT-15, MDT-23 and MDT-24 that control lipid metabolism, stress regulation, muscle development, and responses to pathogen infection (Goh et al., 2014; Grants et al., 2015; Meissner et al., 2009; Nicholas and Hodgkin, 2004; Taubert et al., 2006).

**Figure 1.**
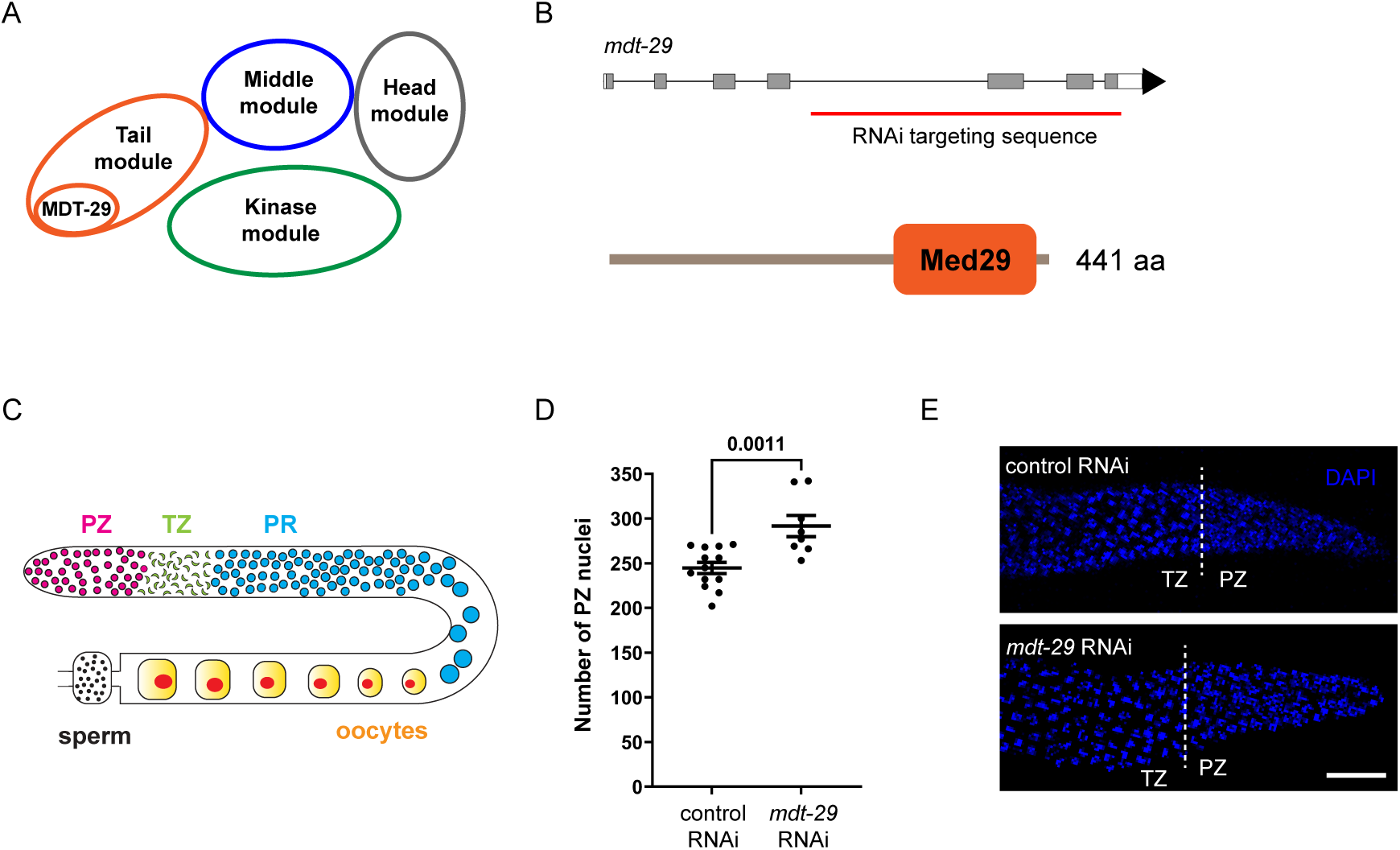
*mdt-29* RNAi knockdown increases germ cell number in the distal progenitor zone. (A) Schematic of the Mediator complex structure showing the head (grey), middle (blue), tail (orange), and kinase modules (green). The MDT-29 subunit is shown within the tail module. (B) Genomic locus (top - grey boxes = exons; black lines = introns; white boxes = untranslated regions), and protein domain structure (bottom) of MDT-29. Med29 = Mediator complex subunit 29 domain. Red line = targeting sequence in RNAi experiments. (C) Schematic of the *C. elegans* adult hermaphrodite germ line. The germ line contains the progenitor zone (∼65% proliferative cells/∼35% cells in the meiotic S-phase), the transition zone (meiotic prophase cells) and the pachytene region (late meiotic cells), followed by mature gametes (oocytes and sperm). (D-E) Quantification (D) and immunofluorescence images (E) of germ cell nuclei number of in the PZ of *rrf-1(pk1417)* one-day adults following *mdt-29* RNAi knockdown. RNAi treatment commenced in L1 larvae and germ lines of one-day adults were examined. n = 13, 8. *p* value assessed by unpaired t-test. Data expressed as mean ± SEM.

Here, we determined the expression and function of MDT-29 in the *C. elegans* germ line, a previously uncharacterized Mediator tail module component. We found that endogenously-tagged MDT-29 is likely expressed in all cell nuclei. Furthermore, MDT-29 protein is concentrated in distinct foci, which resemble phase-separated condensates previously discovered to house Mediator components in mammalian cells (Cho et al., 2018; Sabari et al., 2018). We discovered that germline-specific MDT-29 depletion via RNAi knockdown or auxin-inducible degradation increased cell number within the germline progenitor zone, which is likely driven independently of the GLP-1/Notch signalling pathway. Further analysis suggested that increased cell number in the PZ caused by MDT-29 depletion is the result of an enlarged germline stem cell pool. In addition, we discovered that MDT-29 limits physiological apoptosis in oogenic germ cells. Together, these proliferative and cell death phenotypes likely combine to cause the elevated brood size observed in MDT-29-depleted hermaphrodites. Thus, MDT-29 may coordinate specific gene expression programs to govern germ cell proliferation and death to optimize brood size under certain environmental and/or physiological conditions.

## RESULTS AND DISCUSSION

### MDT-29 is required for *C. elegans* germ cell development

In an ongoing RNA-mediated interference (RNAi) screen for factors that control germ cell development in *C. elegans*, we identified the uncharacterized gene *mdt-29*. *mdt-29* encodes a protein containing a MED29 (Mediator complex subunit 29) domain that is a predicted tail module component of the Mediator complex (Figure 1A-B). Mammalian MDT-29 orthologs (MED29) have been shown to enhance or inhibit cancer cell proliferation depending on the microenvironment (Huang et al., 2024; Kuuselo et al., 2011; Yang et al., 2022).

We assessed MDT-29 function using *mdt-29* RNAi knockdown in the *rrf-1(pk1417)* strain, which permits germline and limited somatic RNAi activity (Figure 1C-E) (Watts et al., 2020; Zou et al., 2019). We found that *mdt-29* knockdown from the L1 stage significantly increased nuclei number in the distal progenitor zone (PZ), proposing an inhibitory function of MDT- 29 in germ cell proliferation (Figure 1D-E). Previous structural analysis of the mammalian Mediator complex suggests that MDT-29/MED29, as part of the tail module, serves as a docking site for transcription factor binding (Tsai et al., 2014). Thus, MDT-29 may suppress germ cell proliferation via transcriptional control of specific signalling pathways. To enable tissue-specific exploration of MDT-29 function, we utilized CRISPR-Cas9 to insert a *degron::green fluorescent protein (gfp)* coding sequence into the 5’ end of the *mdt-29* gene in wild-type animals (Figure 2A) (Dokshin et al., 2018). The degron sequence enables rapid depletion of endogenous MDT-29 protein using the auxin-inducible degradation system (AID) (Zhang et al., 2015). Compared to wild-type animals, insertion of degron::GFP to MDT-29 did not cause overt germline defects, as shown by germ cell nuclei counts and brood size (Figure S1). Thus, the *degron::gfp::mdt-29* strain is a reliable tool for subsequent MDT-29 expression and functional analysis.

**Figure 2.**
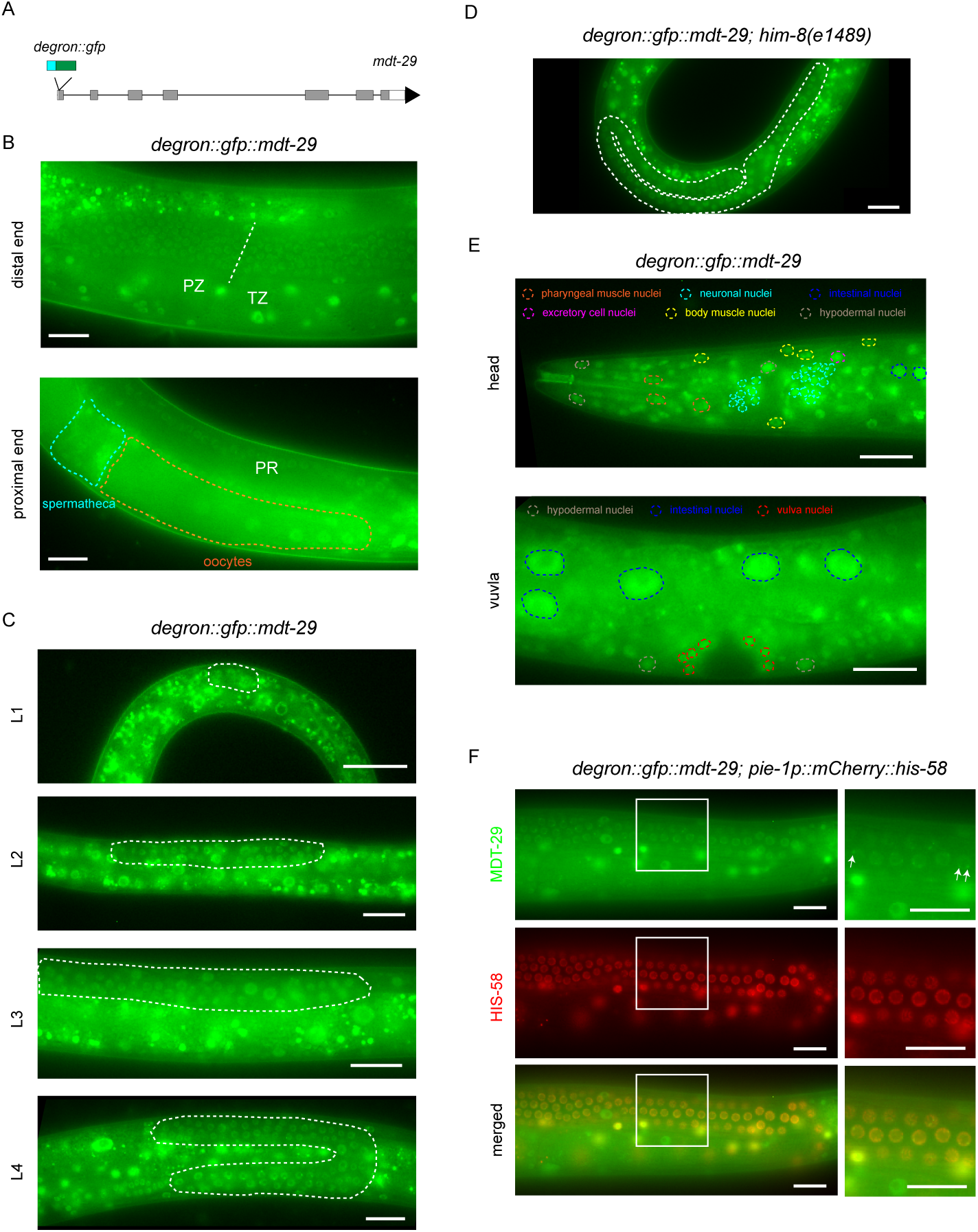
MDT-29 expression pattern analysis. (A) *mdt-29* genomic structure showing insertion of the *degron::gfp* coding sequence at the N-terminus. (B) Fluorescence micrographs of adult germ lines expressing degron::GFP::MDT-29. Expression is detected in the distal end (progenitor zone - PZ and transition zone - TZ, top image) and proximal end (pachytene region - PR, oocytes and spermatheca, bottom image). White dashed line = border between PZ and TZ. Orange = oocytes. Blue = spermathecum. (C) Fluorescence micrographs of larval germ lines expressing degron::GFP::MDT-29. Expression detected at the L1, L2, L3 and L4 stages. White dash = germ line. (D) Fluorescence micrograph of a male adult germ line expressing degron::GFP::MDT-29 in *him-8(e1489)* animals. White dash = germ line. (E) Fluorescence micrographs of somatic tissues expressing degron::GFP::MDT-29. Expression in the head (top) and vulval region (bottom) at the L4 stage. Orange dash = pharyngeal muscle nuclei. Yellow dash = body wall muscle nuclei. Cyan dash = neuronal nuclei. Grey dash = hypodermal nuclei. Blue dash = intestinal nuclei. Magenta dash = excretory cell nucleus. Red dash = vulval nuclei. (F) Co-localisation of degron::GFP::MDT-29 and mCherry::HIS-58 in the adult germ line. White arrows indicate degron::GFP::MDT-29 foci. Green = MDT-29. Red = HIS-58. White box = zoomed region to the right of each image.

### MDT-29 is ubiquitously expressed in *C. elegans*

We next examined the expression pattern of MDT-29 in the *degron::gfp::mdt-29* strain. The Mediator complex functions as an important coregulator of polymerase II-mediated transcription, and thus is widely expressed in *C. elegans* (Moghal and Sternberg, 2003; Steimel et al., 2013; Taubert et al., 2006; Zhang and Emmons, 2001). Based on previous RNA-seq data, MDT-29 is expressed in the soma and germ line (Ebbing et al., 2018; Ghaddar et al., 2023). As *mdt-29* RNAi caused overt germline defects (Figure 1D), we first examined degron::GFP::MDT-29 expression in the germ line. In adult hermaphrodites, degron::GFP::MDT-29 expression was detected in all germ cells, including proliferative germ cells (progenitor zone - PZ), meiotic cells (transition zone - TZ and pachytene region - PR), and in gametes (oocytes and sperm) (Figure 2B). Further, we detected degron::GFP::MDT-29 expression from early embryogenesis and in germ cells throughout larval development (Figures 2C and S2). Previous transcriptomic analysis revealed enrichment of *mdt-29* RNA in the male germ line (Ebbing et al., 2018). To examine MDT-29 expression in males we crossed the *degron::gfp::mdt-29* animals into the *him-8(e1489)* strain. We detected degron::GFP::MDT-29 expression throughout the male germ line (Figure 2D). We also detected degron::GFP::MDT-29 expression in somatic tissues (pharynx, hypodermis, body wall muscle, intestine, nervous system, vulva and the excretory system), consistent with RNA-seq data (Figure 2E) (Ghaddar et al., 2023). Taken together, these observations show that MDT-29 is likely expressed in all tissues throughout *C. elegans* development in both sexes.

### MDT-29 forms condensed foci

While exploring degron::GFP::MDT-29 expression in the germ line, we observed condensed fluorescent foci within germ cell nuclei, suggesting a specific functional locale for MDT-29 (Figure 2F). These degron::GFP::MDT-29 foci were also observed in somatic tissues and throughout animal development (Figure 2E). Such foci are reminiscent of phase-separated condensates previously shown to contain Mediator proteins and RNA polymerase II in mammalian cell models (Cho et al., 2018; Sabari et al., 2018). As mammalian mediator condensates associate with chromatin (Cho et al., 2018; Sabari et al., 2018), we assessed this in *C. elegans*. We found that the chromatin reporter mCherry::HIS-58 and degron::GFP::MDT-29 foci overlap (Figure 2F). These data imply that, as in mammals, the *C. elegans* Mediator complex localizes and concentrates at discrete nuclear locations.

### MDT-29 depletion increases proliferative germ cell number

Our initial RNAi analysis showed that *mdt-29* RNAi knockdown increased germ cell number in the PZ (Figure 1D). However, RNAi-induced mRNA degradation rarely results in complete gene knockdown and the *rrf-1* mutant strain we used is not absolutely specific for germline RNAi (Watts et al., 2020). We thus turned to the tissue-specific auxin-inducible system to confirm MDT-29 germline-specific function. To validate the efficacy of the *degron::gfp::mdt-29* allele for tissue-specific functional analysis, we introduced a transgene (*sun- 1p::TIR1::mRuby*) to express the transport inhibitor response 1 (TIR1) F-box protein (required for AID) in the germ line (Zhang et al., 2015). This new strain (*degron::gfp::mdt- 29; sun-1p::TIR1::mRuby*) exhibits wild-type germline development in the absence of auxin (Figure S1). We found that within 1h of auxin exposure degron::GFP::MDT-29 was efficiently depleted in the adult germ line (Figure 3A). Therefore, degron::GFP::MDT-29 can be rapidly depleted using the AID system.

**Figure 3.**
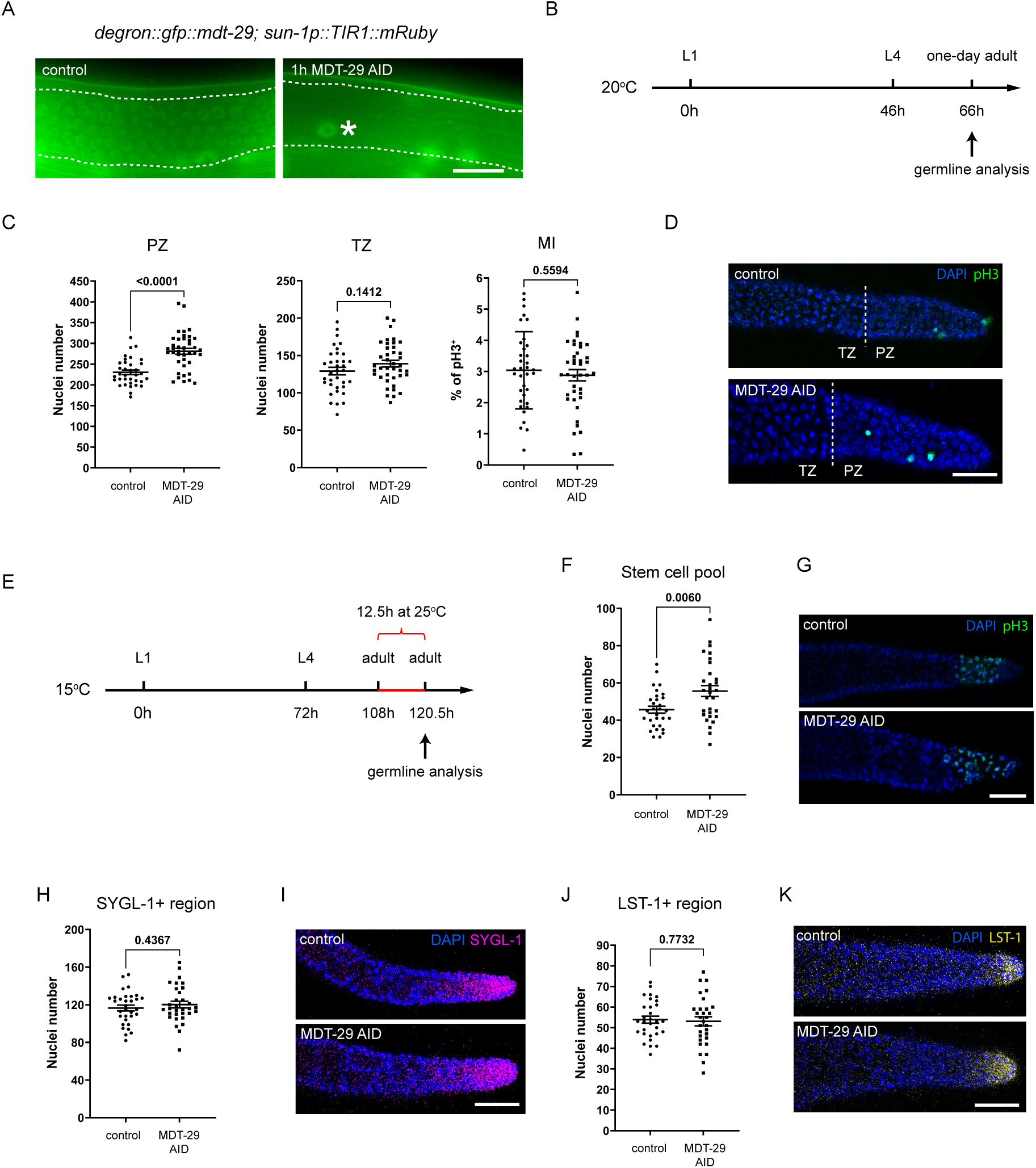
MDT-29 regulation of distal germ cell proliferation. (A) Fluorescence micrographs of *degron::gfp::mdt-29; sun-1p::TIR1::mRuby* animals treated with ethanol (control) or auxin for 1 h. *sun-1p::TIR1::mRuby* = germline-specific AID strain. White dashed line = germ line region. The large GFP positive nucleus (white asterisk) within the dashed line in the 1h MDT-29 AID image is an intestinal nucleus. (B) Timeline of germline analysis following L1 auxin treatment. Freshly hatched L1 animals were transferred to ethanol or auxin plates, incubated at 20°C for 66 hours and germlines dissected in one-day adults. Animals were confirmed as L4 larvae at 46h. (C-D) Quantification (C) and immunofluorescence images (D) of germ cell number of PZ and TZ, and mitotic index (MI) of *degron::gfp::mdt-29; sun-1p::TIR1::mRuby* one-day adults treated with ethanol (control) or auxin for 1 h. Germ lines were stained with DAPI to visualize DNA (blue) and anti-phospho-histone H3 (pH3) to visualize M-phase chromosomes (green). White dashed line = border between PZ and TZ. n = 35, 40. (E) Timeline of stem cell pool evaluation following MDT-29 AID from the L1 stage. Freshly hatched L1 animals were transferred to ethanol or auxin plates, incubated at 15°C for 108h and these one-day adults were heat shocked at 25°C for 12.5h, then their germ lines were dissected and stained at 120.5h. Animals were confirmed as L4 larvae at 72h. (F-G) Quantification (F) and immunofluorescence images (G) of germline stem cells of *emb- 30(tn377); degron::gfp::mdt-29; sun-1p::TIR1::mRuby* one-day adults. Germ lines were stained with DAPI to visualize DNA (blue) and anti-phospho-histone H3 (pH3) to visualize M-phase chromosomes (green). n = 30. (H-I) Quantification (H) and immunofluorescence images (I) of SYGL-1^+^ germ cells of *3xflag::sygl-1*;*degron::gfp::mdt-29;sun-1p::TIR1::mRuby* one-day adults. Germ lines were stained with DAPI to visualize DNA (blue) and anti-FLAG antibodies to visualize SYGL-1^+^ germ cells (pink). n = 32. (J-K) Quantification (J) and immunofluorescence image (K) of LST-1^+^ cells of *3xflag::lst- 1;degron::gfp::mdt-29;sun-1p::TIR1::mRuby* one-day adults. Germ lines were stained with DAPI to visualize DNA (blue) and anti-FLAG antibodies to visualize LST-1^+^ germ cells (yellow). n = 30. Ethanol was applied in the control group. MDT-29 AID commenced from the L1 stage. Data expressed as mean ± SEM. *p* values assessed by unpaired t-test (C, H and J) and Welch’s t-test (F). Scale bars = 20 μm.

Next, we exposed *degron::gfp::mdt-29; sun-1p::TIR1::mRuby* animals to auxin continuously from the L1 larval stage (called MDT-29 L1 AID from here) and analysed the germ lines of one-day old adults (Figure 3B-D). We found that MDT-29 L1 AID caused an increase in the PZ size, consistent with our findings of *mdt-29* RNAi (Figures 1D and 3C-D). However, we did not detect a change in cell number within the TZ when cells are entering early meiotic prophase (Figure 3C-D). The increase in PZ cell number suggested that MDT-29 controls germ cell proliferation.

To evaluate the proliferative state of germ cells following MDT-29 L1 AID, we used an antibody against phospho-histone H3 (pH3) to mark dividing germ cells in M-phase (Figure 3D). The mitotic index (MI) was assessed by calculating the percentage of PZ cells that are at M-phase to signify the mitotic cell cycling activity (Crittenden et al., 2023). We found no significant difference in the MI following MDT-29 L1 AID, suggesting that MDT-29 does not regulate mitotic cell cycling (Figure 3C-D). Together, our RNAi and AID depletion approaches (Figures 1D and 3C) reveal that MDT-29 controls PZ germ cell number in *C. elegans*.

### MDT-29 regulates the distal germline stem cell pool

The germline PZ accommodates ∼65% proliferative cells and ∼35% cells in meiotic S-phase (Fox et al., 2011). Germ cell proliferation requires the maintenance of the germline stem cell pool at the distal end. Germline stem cell renewal is stimulated by the interaction of the GLP-1 Notch receptor expressed on their surface and Notch ligands (LAG-2/APX-1) expressed on somatic distal tip cells (Austin and Kimble, 1987; Gao and Kimble, 1995; Henderson et al., 1994). GLP-1/Notch signalling promotes germline stem cell proliferation through two redundant transcriptional regulators, SYGL-1 and LST-1 (Kershner et al., 2014; Lee et al., 2016; Shin et al., 2017a).

The germline stem cell pool can be estimated using the *emb-30* assay (Cinquin et al., 2010). In the *emb-30(tn377)* temperature-sensitive mutant, the metaphase-to-anaphase transition is blocked at the restrictive temperature (25°C). Therefore, arrested mitotic germ cells, visualized by pH3 immunostaining, represent the entire germline stem cell pool. To assess the role of MDT-29 in germline stem cell pool maintenance, we introduced the *emb-30(tn377)* mutant into the MDT-29 AID strain. *emb-30(tn377);* MDT-29 AID L1 animals were exposed to auxin at 15°C for 106 hours, and then shifted to the restrictive temperature of 25°C for 12.5 hours as adults (Figure 3E). We found that MDT-29 L1 AID increased the germline stem cell pool size by ∼20% (Figure 3F-G). Importantly, auxin exposure alone did not affect the maintenance of germline stem cell pool in *emb-30(tn377)* animals (Figure S3). These results suggest that the increased PZ size induced by MDT-29 L1 AID likely occurs through regulation of the germline stem cell pool.

MDT-29 was previously identified as a putative transcriptional co-activator in the LIN- 12/Notch signalling pathway to facilitate the formation of a transient nuclear complex (Chen et al., 2004). Therefore, we hypothesized that MDT-29 also acts as a co-activator in the related GLP-1/Notch signalling pathway. To investigate this hypothesis, we analyzed the expression of the GLP-1/Notch targets SYGL-1 and LST-1 in the germ line. To this end, we individually crossed endogenously-tagged SYGL-1 (*3xflag::sygl-1*) and LST-1 *(3xflag::lst-1*) animals with the MDT-29 AID strain (Shin et al., 2017a), and counted the number of germ cells in the SYGL-1^+^/LST-1^+^ expression domains (Chen et al., 2020; Shin et al., 2017b). However, we found that MDT-29 L1 AID does not affect the number of SYGL-1^+^ or LST-1^+^ germ cells (Figure 3H-K). These data suggest that the increase in the germline stem cell pool size following MDT-29 L1 AID does not result from altered GLP-1/Notch signalling.

### MDT-29 regulates fecundity

In *C. elegans* hermaphrodites, as germ cells move proximally, mitotic germ cells transit through meiosis and develop into sperm during the L4 larval stage before switching to the oogenesis program in adulthood (Kulkarni et al., 2012). Following meiotic maturation, oocytes are fertilized by sperm and then commence embryogenesis. As MDT-29 is required for proper germ cell proliferation, we examined whether the increase in PZ cell number following MDT-29 AID leads to a difference in progeny output. We found that MDT-29 L1 AID indeed increases the brood size (Figure 4A). We next examined MDT-29 function during gametogenesis, by performing MDT-29 AID from L4, when spermatogenesis initiates. We found that MDT-29 L4 AID did not affect PZ cell number or brood size (Figure 4B-C). These results suggest that MDT-29 is not essential for late germline development and likely acts earlier during larval development to regulate germ cell proliferation.

**Figure 4.**
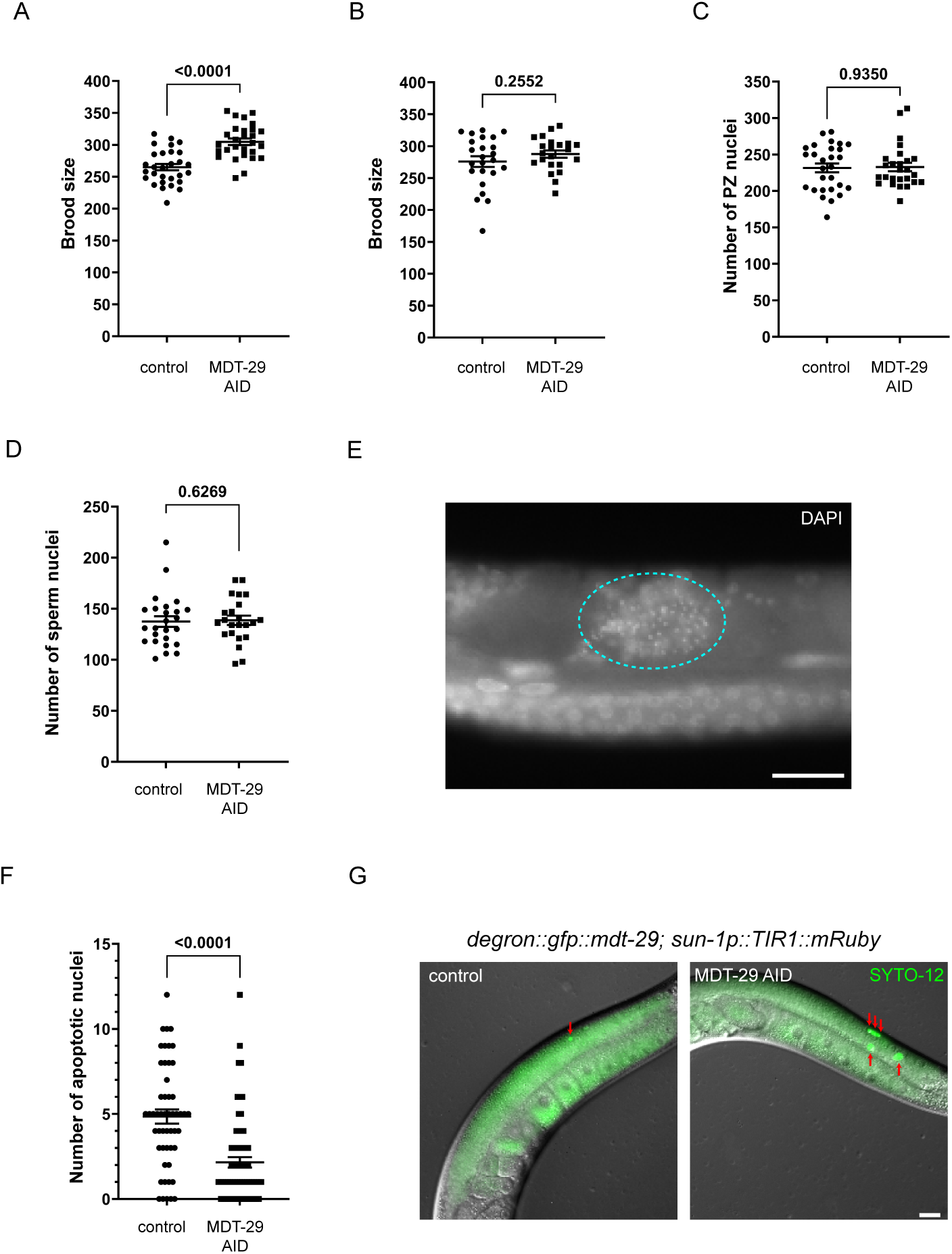
MDT-29 regulation of progeny production. (A) Brood size quantification of *degron::gfp::mdt-29;sun-1p::TIR1::mRuby* hermaphrodites following MDT-29 L1 AID. n=29, 28. (B) Brood size quantification of *degron::gfp::mdt-29; sun-1p::TIR1::mRuby* hermaphrodites following L4 AID. n=22, 24. (C) Quantification of PZ germ cell number of *degron::gfp::mdt-29; sun-1p::TIR1::mRuby* one-day adults following AID treatment from the L4 stage. n = 26, 28 (D-E) Quantification (D) and immunofluorescence image (E) of sperm of *degron::gfp::mdt-29; sun-1p::TIR1::mRuby* one-day adults. Cyan dash = spermathecum. n = 25, 23. (F-G) Quantification (F) and fluorescence micrographs (G) of apoptotic germ cells. SYTO-12 staining was performed following MDT-29 L1 AID. n = 52, 65. Red arrows = apoptotic germ cells. Ethanol was applied in the control group. Data expressed as mean ± SEM. *p* values assessed by unpaired t-test (A-B) and Mann Whitney test (C, D and F). Scale bar = 20 μm.

Because sperm number is a limiting factor in brood size, we investigated whether MDT-29 AID affects the number of sperm. We quantified sperm number of MDT-29 L1 AID one-day adults following germline dissection and DAPI staining, and found that MDT-29 L1 AID did not affect sperm number (Figure 4D-E). During meiotic development, half of the oogenic germ cells undergo physiological apoptosis around the germline loop to supply cytoplasmic components for maturing oocytes (Gumienny et al., 1999). Thus, the number of progeny generated can also be affected by apoptosis. We thus labelled apoptotic germ cells using SYTO-12 following MDT-29 L1 AID, as previously described (Figure 4F-G) (Gumienny et al., 1999). We found that MDT-29 L1 AID reduced the number of apoptotic cells in the pachytene region. Therefore, reduced germ cell apoptosis may contribute to the increase in brood size observed when MDT-29 is depleted in the germ line.

Taken together, our study reveals that MDT-29, a tail module Mediator complex component, limits fecundity in *C. elegans*. Our analysis of germ cell behavior suggests that MDT-29 controls progeny production by modulating the germline stem cell pool size and germ cell apoptosis. Thus, MDT-29 likely controls discrete gene regulatory networks to control germ cell behavior by coordinating with context-specific TFs.

### Methods Strains

*C. elegans* strains were maintained on Nematode Growth Medium (NGM) plates seeded with OP50 *Escherichia coli* bacteria at 20°C, unless otherwise stated. Animals were continuously fed for at least three generations prior to analysis. All strains used in this study are listed in Table S1.

### Endogenous tagging with CRISPR-Cas9

The *degron::GFP::mdt-29* allele was generated using CRISPR-Cas9 genome editing (Dokshin et al., 2018). The crRNA targeting *mdt-29* was designed and ordered using the online tool provided by https://sg.idtdna.com (Table S2). Ultramer primers were used to produce asymmetric-hybrid donor repair templates (Table S2). Next, the injection mixture was introduced into wild-type animals, which contained 2 μg universal tracrRNA, 1.1 μg crRNA, 5 μg Cas9 protein, 4 μg repair template, and *myo-2::mCherry* plasmid (4ng/μL). F1 progeny of injected wild-type animals were individually isolated, and their progeny were screened for degron::GFP insertion using PCR. The insertion was verified by Sanger sequencing.

#### RNAi experiments

*mdt-29* RNAi was performed by the standard feeding protocol (Fraser et al., 2000). The *mdt-29* RNAi plasmid was obtained from the Vidal RNAi library (Rual et al., 2004). The *mdt-29* and L4440 (empty vector control) plasmids were transformed into the HT115 *E. coli* strain, and grown in Luria Broth (LB) media with 100ug/μl ampicillin at 37°C for 16 hours. RNAi bacteria was seeded on RNAi plates (NGM plates + 3mM IPTG). Selected animals were maintained on RNAi plates for various periods depending on the experiment (described in Figure legends).

#### Auxin-inducible degradation (AID) experiments

AID experiments were performed by transferring worms to ethanol (control) or auxin plates (Zhang et al., 2015). The 400mM auxin (indole-3- acetic acid, Alfa Aesar, ALFA10556) stock solution was dissolved in ethanol. Auxin plates (1mM) were prepared by diluting auxin stock solution into NGM agar before pouring plates.

#### Brood size analysis

Brood size was quantified by counting progeny produced by each worm in the first 4 days of adulthood. L4 worms were picked onto individual NGM plates, allowing egg-laying for 24 hours. Next, mothers were moved to new plates every 24 hours. The number of hatched larvae were counted. If RNAi knockdown or AID was required, the brood size analysis was conducted on RNAi or auxin plates.

#### Germline staining

Germ lines were isolated from sedated worms before immunostaining. Dissected germ lines were fixed on poly-L-lysine coated slides by immersing in ice-cold methanol for 1 minute then 3.7% paraformaldehyde for 30 minutes. Fixed germ lines were washed twice with PBST (PBS + 0.5% Tween 20) and blocked using 30% normal goat serum. Then germ lines were incubated in primary antibody at 4°C overnight and washed twice with PBST. Next, germ lines were incubated in secondary antibody and DAPI for 2 hours at 20°C, followed by two PBST washes. Samples were mounted on slides using Fluoroshield mounting media (Sigma). Stained germ lines were imaged using confocal microscopes Leica SP5 (63x objectives). Primary antibodies used: anti-phospho-Histone H3 (Ser10) (06-570, Merk, 1:500), monoclonal ANTI-FLAG® M2(F1804, Sigma-Aldrich, 1:500). Secondary antibodies used: goat anti-rabbit 647 (A21245, Invitrogen, 1:1000), goat anti-mouse 555 (A21422, Invitrogen, 1:1000).

#### Germline analysis

Germline analysis was performed as previously described (Gopal et al., 2017). Z-stack images were acquired using a Leica SP5 confocal microscope (63x objectives), and reconstructed into 3-D germline models by using Imaris Suite 9 software. Nuclei of mitotic germ cells (PZ) and pachytene germ cells (PR) exhibit globular morphology. The TZ, which accommodates germ cells at leptotene and zygotene stages, was defined as a continuous region where germ cell rows contain over 60% of crescent shape nuclei. The germline region before the TZ is defined as the PZ. The germline region after the TZ but before the germline loop is defined as the pachytene region. Semi-automatic nuclei counting was performed using the Imaris Suite 9 to count germ cell nuclei in each region, as in our previous study (Gopal et al., 2017). The nuclei diameter of PZ is set as 2.4 μm, and the nuclei diameter of TZ is set as 2.4 μm in width and 1.8 μm in height. The reliability of this method is confirmed as the readout of PZ nuclei number in control samples is within the previously reported range (Gopal et al., 2017).

#### Fluorescence microscopy

Worms were anesthetised in 0.01% tetramisole and mounted on a 5% agarose pad. Images were taken using an Axio Imager M2 fluorescence microscope and analysed using Zen software (Zeiss).

#### Analysis of apoptotic germ cells

Apoptotic germ cells were analysed using SYTO-12 staining. One-day old adult worms were washed using M9 media and transferred into a prepared 50 μM SYTO-12 solution (Thermo Scientific™) containing OP50 bacteria. After incubating at 25°C for 5 hours, worms were transferred to freshly seeded NGM plates for 1 hour, allowing the expulsion of stained bacteria from the gut. The number of apoptotic germ cells within the germ line (SYTO-12^+^) was counted using fluorescence microscopy.

#### Estimation of stem cell pool

The stem cell pool was estimated using the *emb-30* assay as previously described (Cinquin et al., 2010). The *emb-30(tn377); degron::gfp::mdt-29; sun-1p::TIR1::mRuby* strain was maintained at the permissive temperature of 15°C. *emb-30(tn377); degron::gfp::mdt-29; sun-1p::TIR1::mRuby* worms were synchronized at the L4 stage (Christmas-tree substage) and maintained at 15°C for 36 hours to grow into adults. These animals were then switched to the restrictive temperature of 25°C for 12.5 hours, at which the metaphase-to-anaphase transition was blocked (Furuta et al., 2000). Germ lines were then isolated and immunostained using DAPI, anti-phospho-Histone H3 (Ser10) antibody to respectively visualize nuclei and M-phase cells using a Leica SP5 confocal microscope. Germline stem cells (pH3^+^ cells in the PZ) were quantified using the Imaris Suite 9 software.

#### Sperm quantification

Sperm quantification was conducted as previously described (Dufourcq-Sekatcheff et al., 2021). Worms were dissected and fixed on the poly-L-lysine coated slides. After DAPI staining, z-stack images of spermatheca were obtained using the AxioImager M2 (Zeiss) fluorescence microscope (40x objective). The number of sperm was manually counted using the FIJI (Fiji Is Just) ImageJ 2.14.0/1.54f software.

#### Statistical analysis

Statistical analyses were conducted in GraphPad Prism 10 using two-tailed unpaired t-tests or one-way analysis of variance (ANOVA) test. In unpaired t-tests, Welch’s correction was applied when significant variance was observed, and Mann-Whitney test was used when data were not normally distributed. Ordinary two-tailed ANOVA test was performed when data were normally distributed, otherwise, Kruskall–Wallis test was performed. Specific statistical tests used for each experiment are detailed in the Figure legends. Data are expressed as mean ± S.E.M. Significant differences was determined with a p value < 0.05.

## Acknowledgements

We thank members of the Pocock for advice and comments on the manuscript. Imaging was performed at Monash Microimaging. Some strains were provided by the Caenorhabditis Genetics Center (University of Minnesota), which is funded by the NIH Office of Research Infrastructure Programs (P40 OD010440).

## Author Contributions

Conceptualization: WC, RP

Methodology: QF, CT, WC, RP

Investigation: QF, CT, WC, RP

Visualization: QF, CT, WC, RP

Funding acquisition: QF, WC, RP

Project administration: QF, CT, WC, RP

Supervision: WC, RP

Writing – original draft: QF

Writing – review & editing: QF, CT, WC, RP

## Data and materials availability

All data is available in the main text, supplementary materials, and source data. No accession codes, unique identifiers, or weblinks are in our study and there are no restrictions on data availability. Materials are available upon request from Roger Pocock.

## Supplementary data

**Figure S1.**
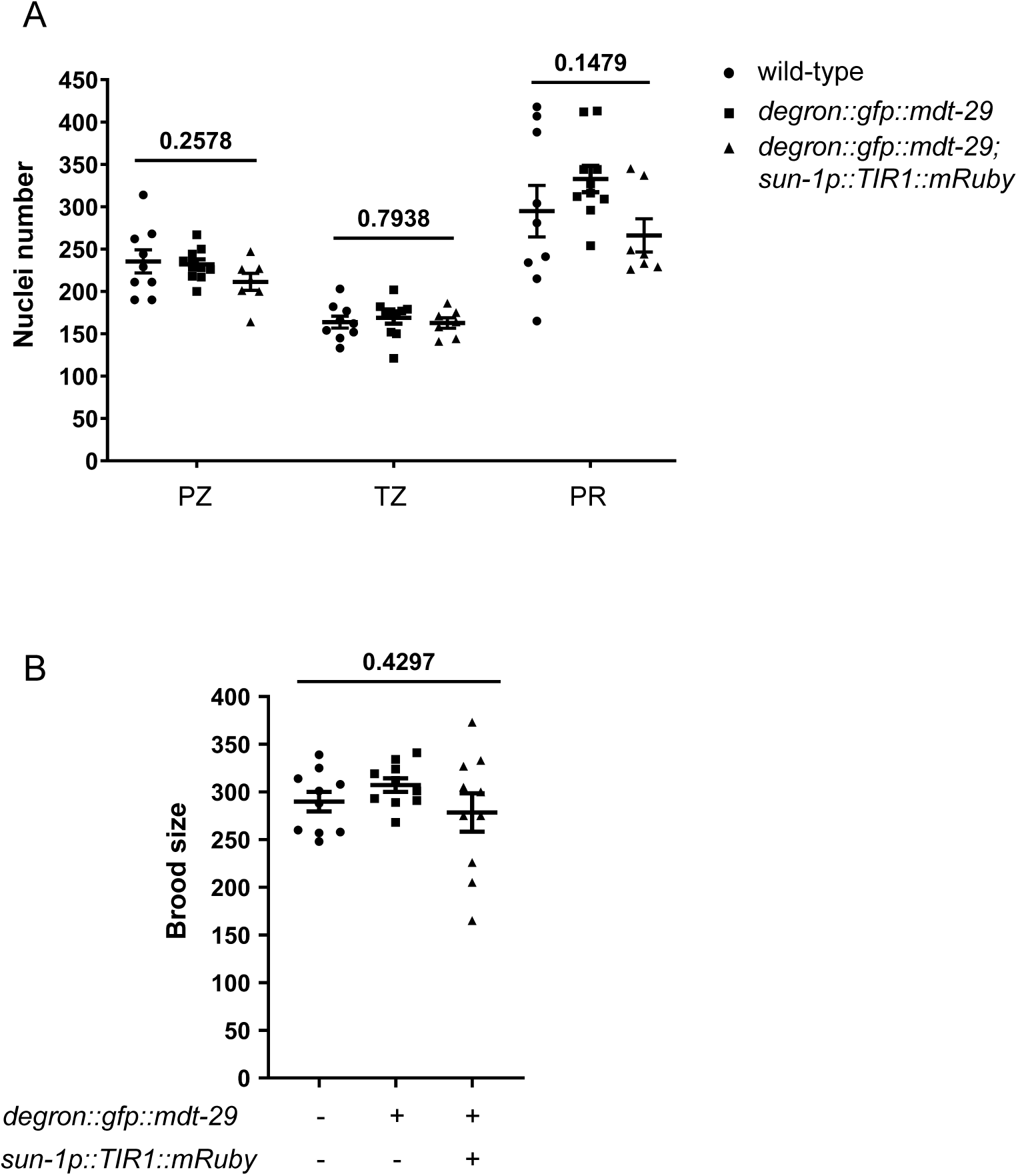
Tagging of MDT-29 with degron-GFP does not affect germline function. (A) Germ cell number quantification of the PZ, TZ and PR of wild-type, *degron::gfp::mdt-29* and *degron::gfp::mdt-29; sun-1p::TIR1::mRuby* animals. n = 7, 9,10. *p* values assessed by one-way ANOVA. Data expressed as mean ± SEM. (B) Brood size quantification wild-type, *degron::gfp::mdt-29* and *degron::gfp::mdt-29; sun-1p::TIR1::mRuby* animals. n = 10. *p* value assessed by Kruskal-Wallis ANOVA test. Data expressed as mean ± SEM.

**Figure S2.**
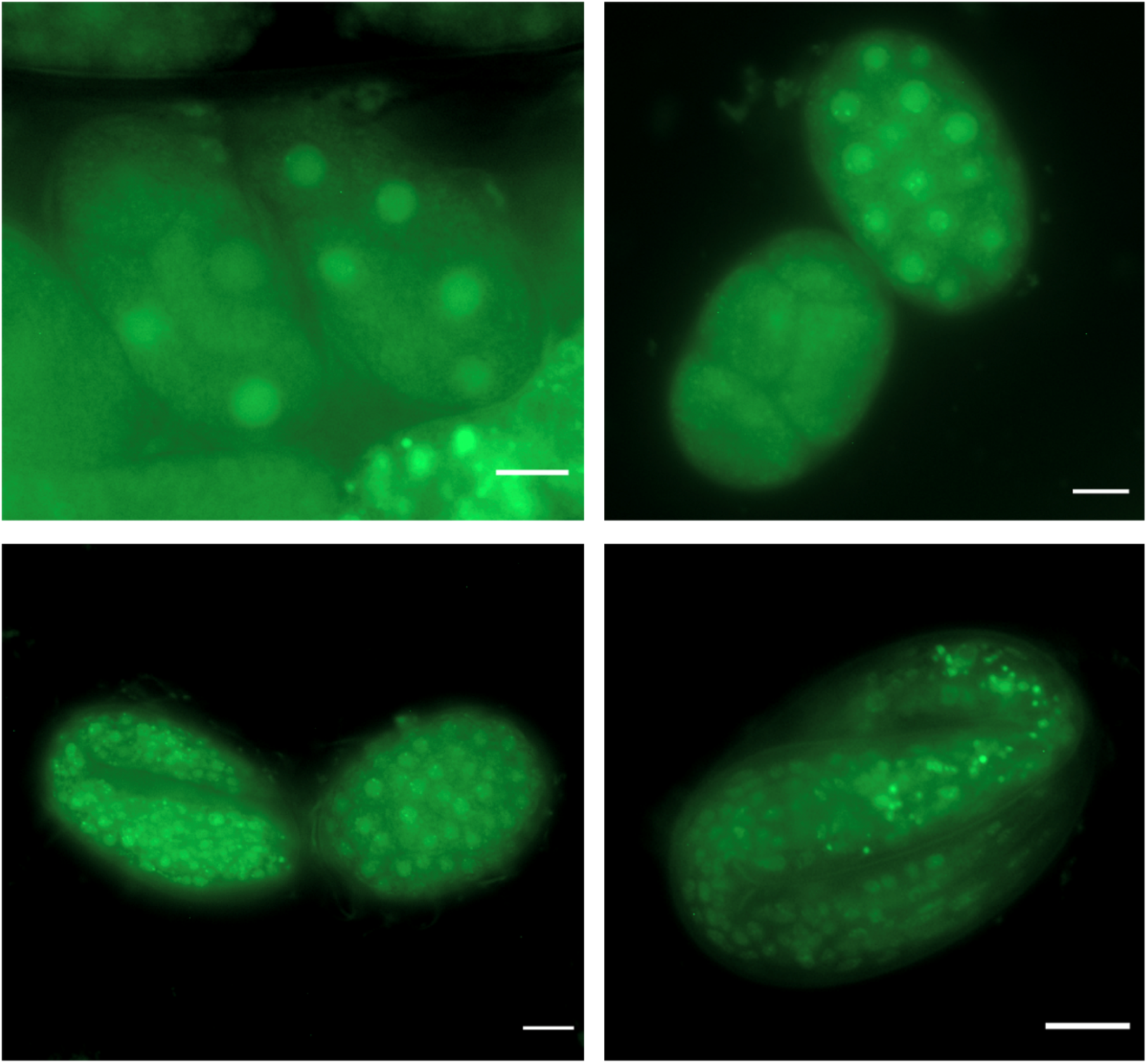
MDT-29 is expressed throughout embryogenesis. Fluorescence images of *degron::gfp::mdt-29* embryos showing nuclear expression from early to late embryogenesis. Scale bars = 10 μm.

**Figure S3.**
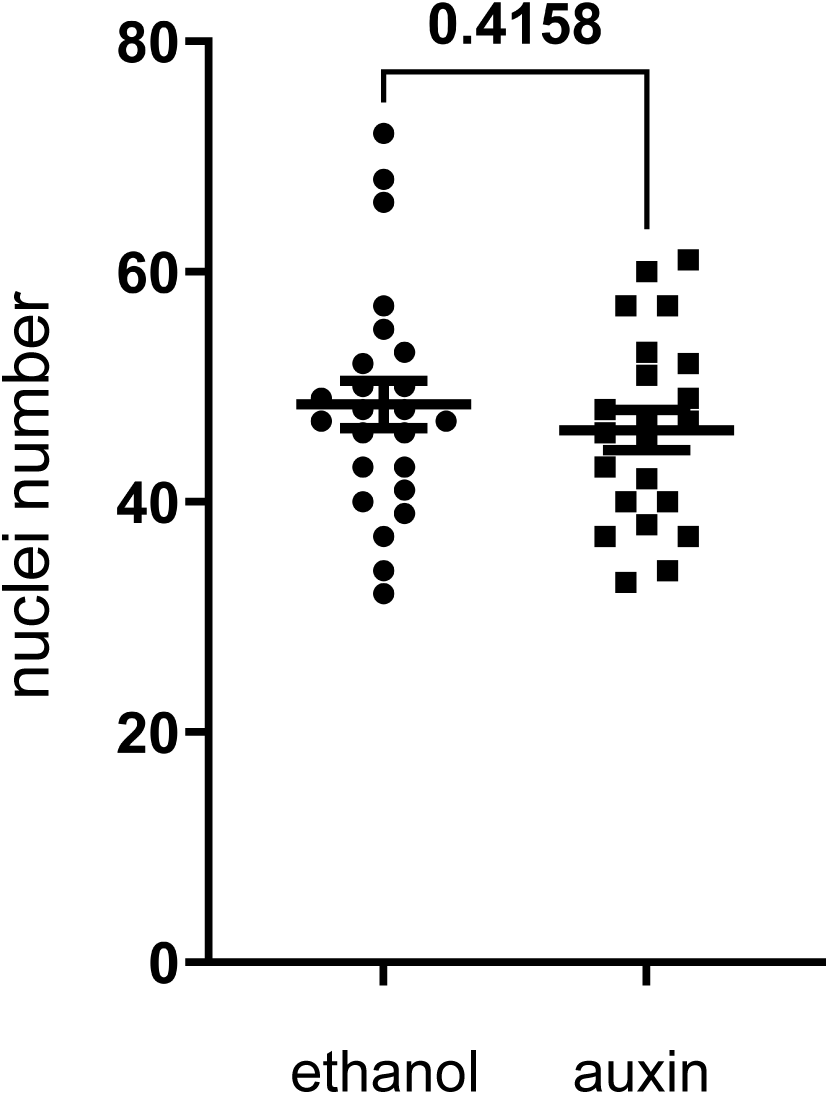
Auxin exposure does not affect the germline stem cell pool in the *emb-30(tn377)* strain. The germline stem cell pool size was evaluated in *emb-30(tn377)* animals +/– auxin treatment from the L1 stage. Data expressed as mean ± SEM. *p* value assessed by unpaired t-test.

